# Tissue growth factor β stimulates fibroblast-like synovial cells to produce extra domain A fibronectin in osteoarthritis

**DOI:** 10.1101/335562

**Authors:** TW Kragstrup, DH Sohn, CM Lepus, K Onuma, Q Wang, WH Robinson, J Sokolove

## Abstract

**Introduction:** The pathophysiology of osteoarthritis (OA) involves wear and tear, and a state of low-grade inflammation. Wear and tear leads to tissue degradation and tissue repair responses, including tissue growth factor beta (TGFβ)-induced myofibroblast production of extracellular matrix (ECM). Fibronectins are an essential part of the ECM, and injection of fibronectin fragments into rabbit joints is a previously established animal model of OA. Alternatively-spliced fibronectin contains the ED-A domain (ED-A FN) and has been shown to activate Toll-like receptor 4. In this study, we tested the hypothesis that FN fragments containing the ED-A domain could be one mechanism transducing mechanical events into inflammatory signals in OA.

**Methods:** Samples of synovial membrane and cartilage were obtained from patients with knee OA undergoing joint replacement surgery. Immunostaining for ED-A FN and the myofibroblast marker alpha smooth muscle actin (αSMA) was performed on synovial membranes and fibroblast-like synovial cells (FLS). FLS were stimulated with TGFβ, TNFα, lipopolysaccharide, IL-6, OA synovial fluid, or chondrocyte lysate, and analyzed for ED-A FN. Synovial cells isolated by enzymatic digestion and human monocyte-derived macrophages (MDM) were incubated with recombinant ED-A FN, plasmin, cellular FN, or cellular FN digested with plasmin; and culture supernatants were analyzed for MCP-1 and TNFα.

**Results:** We hypothesized that ED-A FN is produced by OA FLS in response to factors found in the OA synovial joint. Indeed, the production of ED-A FN by OA FLS was increased by TGFβ, OA synovial fluid, and lysed chondrocytes in all experiments (n=3). ED-A FN co- localized with the myofibroblast marker αSMA in both the OA FLS (n=3) and in the OA synovial membranes (n=8). We further hypothesized that ED-A FN expression is associated with cellular density and expression of inflammatory molecules in OA. ED-A FN staining was associated with both number of lining layer cells (rho=0.85 and p=0.011) and sublining cells (rho=0.88 and p=0.007) in the OA synovium (n=8), and co-localized with both MCP-1 and TNFα (n=5). Recombinant ED-A FN increased the production of both MCP-1 and TNFα by MDM (n=3) and OA FLS (n=3). Finally, we demonstrated that the FN fragments containing the ED-A domain generated the same production of both MCP-1 and TNFα as recombinant ED-A FN.

**Conclusion:** The disease process in OA shares features with the chronic wound healing response including myofibroblast differentiation and production of mediators that promote myofibroblast production of ED-A FN. We show that recombinant and plasmin-derived ED-A fragments stimulate FLS and MDM to produce pro-inflammatory mediators. Our findings support utilizing ED-A FN for drug delivery or therapeutic targeting of the formation of ED- A FN or the enzymatic fragmentation of FN to reduce pro-inflammatory responses in OA.

## Background

Osteoarthritis (OA) is a very common disease affecting approximately 60% of the population by age 65 years with no current therapeutics approved for preventing disease progression. The pathophysiology involves wear and tear and a state of low-grade inflammation, which has generated the hypothesis that OA could be a chronic wound [1-3]. Thus, wear and tear leads to tissue degradation followed by tissue repair responses including tissue growth factor beta (TGFβ) induced myofibroblast production of extracellular matrix (ECM) [4]. ECM molecules and breakdown products can then function as danger associated molecular patterns (DAMPs) leading to activation of the immune system through activation of Toll-like receptors (TLRs) on macrophages [5].

Fibronectins (FNs) are an essential part of the ECM. The active FNs consist of several isoforms and peptide fragments with individual functions [6]. FN produced by fibroblasts is thus a result of both alternative splicing potentially incorporating extra domain A (ED-A) or ED-B and proteolytic cleavage by enzymes such as plasmin [7].

The association between FN and OA is already established [8,9]. Injection of FN fragments in rabbit joints lead to many characteristics of OA including cartilage degradation and bony spur formation and is now an established animal model of OA [10]. FN fragments are present in OA synovial fluid and have been shown to induce pro-inflammatory cytokines and matrix metalloproteinases [11,12]. However, the mechanisms coupling FN fragments and OA are not fully understood. The FN fragments containing the ED-A domain are of particular interest because they show properties with possible implications for both immune activation and joint damage in OA [13]. These fragments have thus recently been shown to function as TLR-4 agonists [14], and synovial fluid levels correlate with establishment of disease and radiographic progression in rheumatoid arthritis [15,16]. Also, ED-A FN is produced by myofibroblasts and are essential for normal wound healing fitting the theory of OA as a chronic wound [17-19].

In this study, we hypothesize that ED-A FN fragments represent a factor transducing mechanical events into inflammatory signals in OA.

## Materials and methods

### Osteoarthritis patients

Samples were obtained from patients with knee OA undergoing joint replacement surgery. Patients were diagnosed with knee OA of Kellgren-Lawrence score 2 to 4 according to the 1985 criteria of the American Rheumatism Association [20,21].

### Ethics, consent and permissions

Samples were obtained under protocols approved by the Stanford University Institutional Review Board and with the patients’ informed consent.

### Isolation of synovial tissue, synovial cells, synovial FLSs and chondrocytes

Samples from the joint replacement surgery included synovial membrane, synovial fluid, and cartilage. Synovial tissue samples for immunofluorescence were snap frozen in Tissue-Tek. Synovial cells for immunofluorescence and in vitro experiments were isolated after enzymatic digestion of the tissue. Synovium was minced with sterile scissors digested with collagenase grade II (Clostridium histolyticum) in DMEM and antibiotics supplemented with 5% fetal bovine serum (FBS). The resulting cell suspension was pipetted through a 70 μm mesh and centrifuged at 250g for 10 min. The supernatant was removed and after washing the cells were incubated at 37°C and 5% CO_2_. Passage 5 cells were used for immunofluorescence of FLSs and passage 1 cells were used for in vitro experiments with synovial cells. Synovial fluid from an OA patient was centrifuged and cell free synovial fluid was frozen at −80°C until use. Chondrocyte lysate was made after isolating chondrocytes from the cartilage of an OA patient. Cartilage was diced and digested for 30 min at 37°C in 1 mg/ml pronase (Streptomyces griseus pronase E) in 10 ml DMEM F12 and antibiotics supplemented with 5% FBS. After washing cells were digested for 18 hours with 3 mg/ml collagenase grade II (Clostridium histolyticum) in 25 ml DMEM F12 and antibiotics supplemented with 5% FBS. The resulting cell suspension was pipetted through a 70 μm mesh, centrifuged at 250g for 10 min and resuspended in DMEM F12. The chondrocytes were lyzed by two cycles of snap freezing cells in −170°C and thawing cells in a 37°C water bath. The resulting protein concentration was 1.7 mg/ml. Human monocyte-derived macrophages (MDMs) were generated from peripheral blood mononuclear cells (PBMCs) by density-gradient centrifugation of LRS chamber content (Stanford Blood Center) over Ficoll (Invitrogen), purification of human monocytes by negative selection (Miltenyi Biotec), and differentiation of the monocytes into macrophages by culturing them for 7 days in RPMI containing 10% FBS and 30 ng/ml of human M-CSF.

### Immunoflourescence of osteoarthritis fibroblast-like synoviocytes

Immunoflourescence of FLSs were done as previously described.[22] Sterile glass slides were placed in 24-well cell culture plates. FLSs were then seeded at a concentration of 5.0×10^4^ cells/mL in DMEM and antibiotics supplemented with 5% FBS and incubated for 24 hours at 37°C. The cells were either untreated or stimulated with TGFβ at 5 ng/ml, TNFα (PeproTech) at 10 ng/mL, lipopolysaccharide (LPS) (Sigma) at 100 ng/ml, IL-6 (PeproTech) at 10 ng/ml, OA synovial fluid at 1%, or chondrocyte lysate at 1%. Cells were fixed with 4% paraformaldehyde and then incubated for 30 min at room temperature with PBS with 0.05% Tween20, 1% bovine serum albumin, 5% normal goat serum, and 0.03% NaN_3_. Cells were stained with rabbit polyclonal anti-EDA FN (kind gift from Digna Bioscience, Madrid, Spain), mouse IgG2a anti-αSMA (clone 1A4, R&D Systems), rabbit polyclonal anti-αSMA (Abcam), mouse IgG2a anti-CD45 (clone F10-89-4, Abcam), mouse IgG1 anti-CD31 (clone JC70A, Dako), mouse IgG2b anti-MCP-1 (clone 23002, R&D Systems), and rabbit polyclonal anti-TNFα (Abcam). Rabbit polyclonal isotype, mouse IgG2a isotype, mouse IgG1, and mouse IgG2b were used as negative controls. Goat anti-rabbit alexa 488, goat anti- mouse IgG2a alexa 555, goat anti mouse IgG1 alexa 647, and goat anti-mouse IgG alexa 555 were used as secondary antibodies (all Thermo Fischer). Wells were washed twice, and glass slides were mounted with Prolong Gold Antifade with DAPI (Thermo Fischer). All micrographs were collected by using a Zeiss LSM-710 confocal microscope. ED-A FN positive cells and total cells were counted for each culture condition.

### Immunoflourescence of osteoarthritis synovial membranes

Synovial tissue samples for immunofluorescence were snap frozen in Tissue-Tek and cut in sections of 10 μm using a microtome and sections were fixed with 4% paraformaldehyde and then incubated for 30 min at room temperature with PBS with 0.05% Tween20, 1% bovine serum albumin, 5% normal goat serum, and 0.03% NaN3. Cells were stained with rabbit polyclonal anti-EDA FN (kind gift from Digna Bioscience, Madrid, Spain) and mouse IgG2a anti-α-smooth muscle actin (αSMA) (clone 1A4, R&D Systems). Rabbit polyclonal isotype and mouse IgG2a isotype were used as negative controls. Goat anti-rabbit alexa 488 and goat anti-mouse IgG2a alexa 555 were used as secondary antibodies (both Thermo Fischer). Wells were washed twice, and glass slides were placed in 2 μL of Prolong Gold Antifade with DAPI (Thermo Fischer) and allowed to dry overnight. All micrographs were collected with a Zeiss LSM-710 confocal microscope. Micrographs were analyzed using ImageJ software (NIH).The area with ED-A staining was found and calculated as percentage of total area. The area with DAPI staining in the sublining layer was found and calculated as percentage of total area. The lining layer cell thickness was counted.

### Stimulation of macrophages and OA synovial cells

For confirmation that endotoxin contamination was not a confounding factor, we stimulated murine RAW 264.7 macrophages with ED-A FN at 10 μg/ml in the presence of 15 μg/ml of polymyxin B (Sigma-Aldrich) or after treatment of ED-A FN with proteinase K (20 μg/ml) followed by boiling for 15 minutes or boiling for 30 minutes only. The monocyte-derived macrophages and OA synovial cells were thawed and cultured in RPMI medium supplemented with 10% FBS, penicillin, streptomycin, and glutamine at a density of 2 x 106 cells/ml. Then, cells were incubated with recombinant ED-A FN (kind gift from Digna Bioscience, Madrid, Spain) at 100 ng/ml, plasmin (Sigma-Aldrich) at 0.5 ng/ml, cellular FN (Sigma-Aldrich) at 1000 ng/ml or cellular FN pre-incubated with plasmin for 30 min at 37°C at the same concentrations. For each experiment, an untreated cell culture with the same number of cells in medium without stimulants was used for comparison and a culture stimulated with LPS at a concentration of 100 ng/ml was used as a positive control. Cells were cultured for 48 h at 37°C in a humidified incubator with 5% CO_2_ without changing of medium. After incubation, supernatants were stored frozen at −80°C for later assessment of MCP-1 and TNFα content by ELISA (both PeproTech).

## Statistics

Statistical analyses were performed using GraphPad Prism 6.0 for Mac (GraphPad Software). Correlations were made using Spearman’s Rho. Ratios were log transformed and analyzed with the paired t-test. In all tests, the level of significance was a two-sided P value of less than 0.05.

## Results

### ED-A FN is produced by OA FLSs in response to TGFβ, OA synovial fluid, or chondrocyte lysate

We first hypothesized that ED-A FN is produced by OA FLSs in response to factors present in the OA synovial joint. To test this hypothesis OA FLSs were incubated with TGFβ, OA synovial fluid and proteins from lysed chondrocytes (n=3). Spontaneous production of ED- A FN was found in a small number of cells in all cultures. The production of ED-A FN by OA FLSs was increased by TGFβ, OA synovial fluid, and lysed chondrocytes in all experiments. There was no change in the production of ED-A FN by OA FLSs when using TNFα, LPS, or IL-6. No signal was detected when staining with negative control isotype antibody or when incubating lysed chondrocytes or OA synovial fluid in wells without OA FLSs (Figure 1A).The ED-A FN staining in the OA FLSs co-localized with staining of the myofibroblast marker αSMA (Supplementary figure 1).

**Figure 1.**
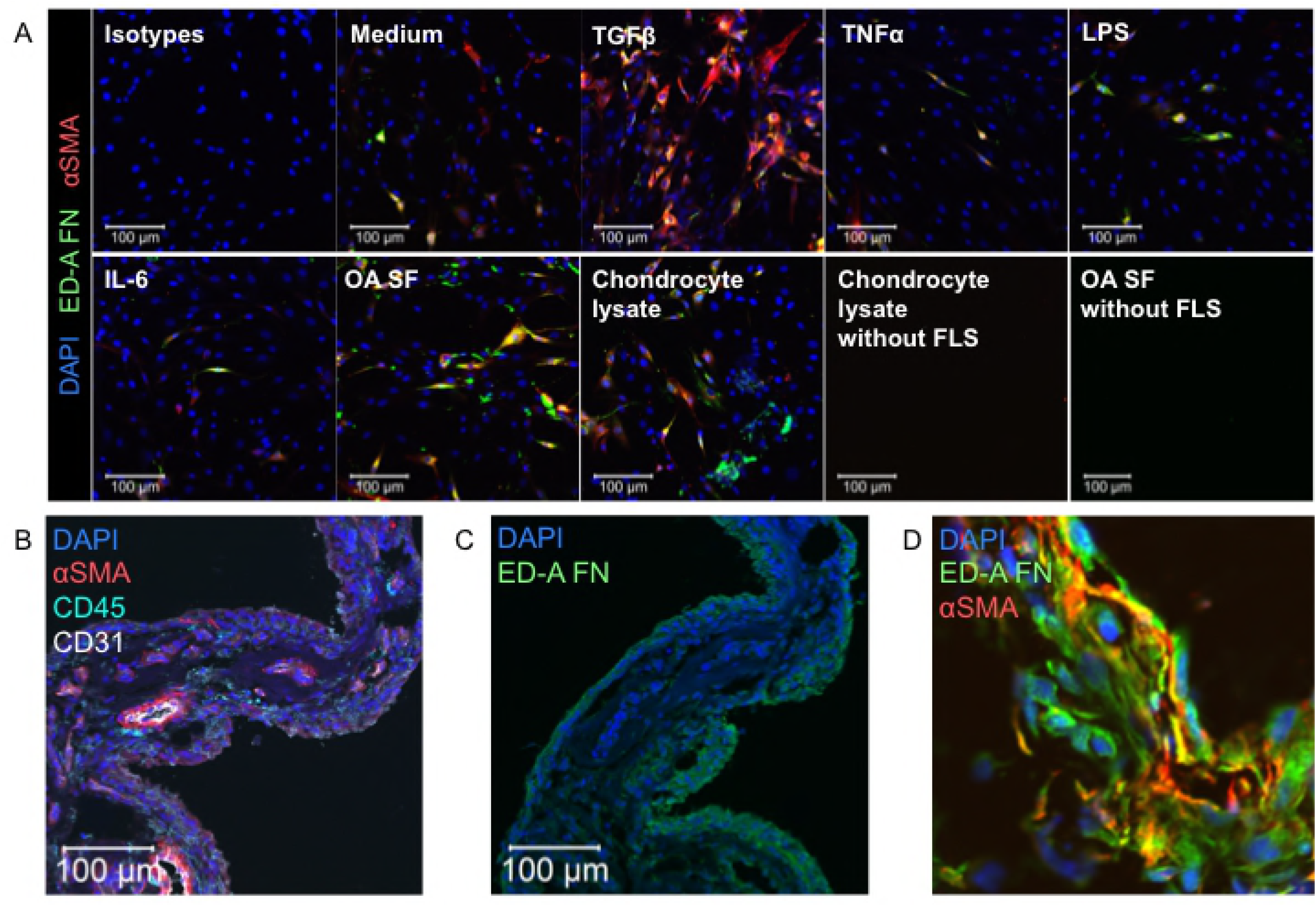
ED-A FN production by OA FLSs and ED-A FN expression in OA synovium. **A**. Representative confocal microscopy images of αSMA (red) and ED-A FN (green) in OA FLS cultures incubated with medium, TGFβ, TNFα, LPS, IL-6, OA SF, and OA chondrocyte lysate (n=3). ED-A FN production by FLSs was increased by adding TGFβ, cell free OA SF, or lysed OA chondrocytes. No staining was seen using isotype control antibodies or adding chondrocyte lysate to a well without FLSs. **B-D**. Representative confocal microscopy images of CD45, CD31, αSMA and ED-A FN in OA synovium (n=8). ED-A FN and αSMA positive cells were found around blood vessels and in the lining layer. **D**. Representative confocal microscopy images of ED-A FN co-localization with αSMA (n=3).

### ED-A FN is located to αSMA^+^ myofibroblasts in OA synovium

We then tested whether ED-A FN also co-localized with αSMA in the OA synovial membrane. Both ED-A FN staining and αSMA staining were found in the synovium from all OA patients (n=8) (Figure 1B-D and Supplementary figure 2). The ED-A FN staining was most intense in lining layer cells while αSMA staining was most intense in cells surrounding CD31 positive blood vessels. However, most ED-A FN positive cells were also to some extent αSMA positive in all the stained synovial membranes (n=3). These results indicate that ED-A containing FN can be produced by OA myofibroblasts in response to mediators involved with tissue regeneration in vitro and in vivo.

### ED-A FN is associated with increased number of lining layer cells and sublining cells in OA synovium

Next, we hypothesized that ED-A FN expression associates with inflammation in OA. Therefore, we analyzed the association between the degree of ED-A FN staining and the number of lining layer cells and sublining cells in OA synovium. The ED-A FN staining associated with both number of lining layer cells (rho=0.85 and *p*=0.011) and sublining cells (rho=0.88 and *p*=0.007) in OA synovium (n=8) (Figure 2A-C).

**Figure 2.**
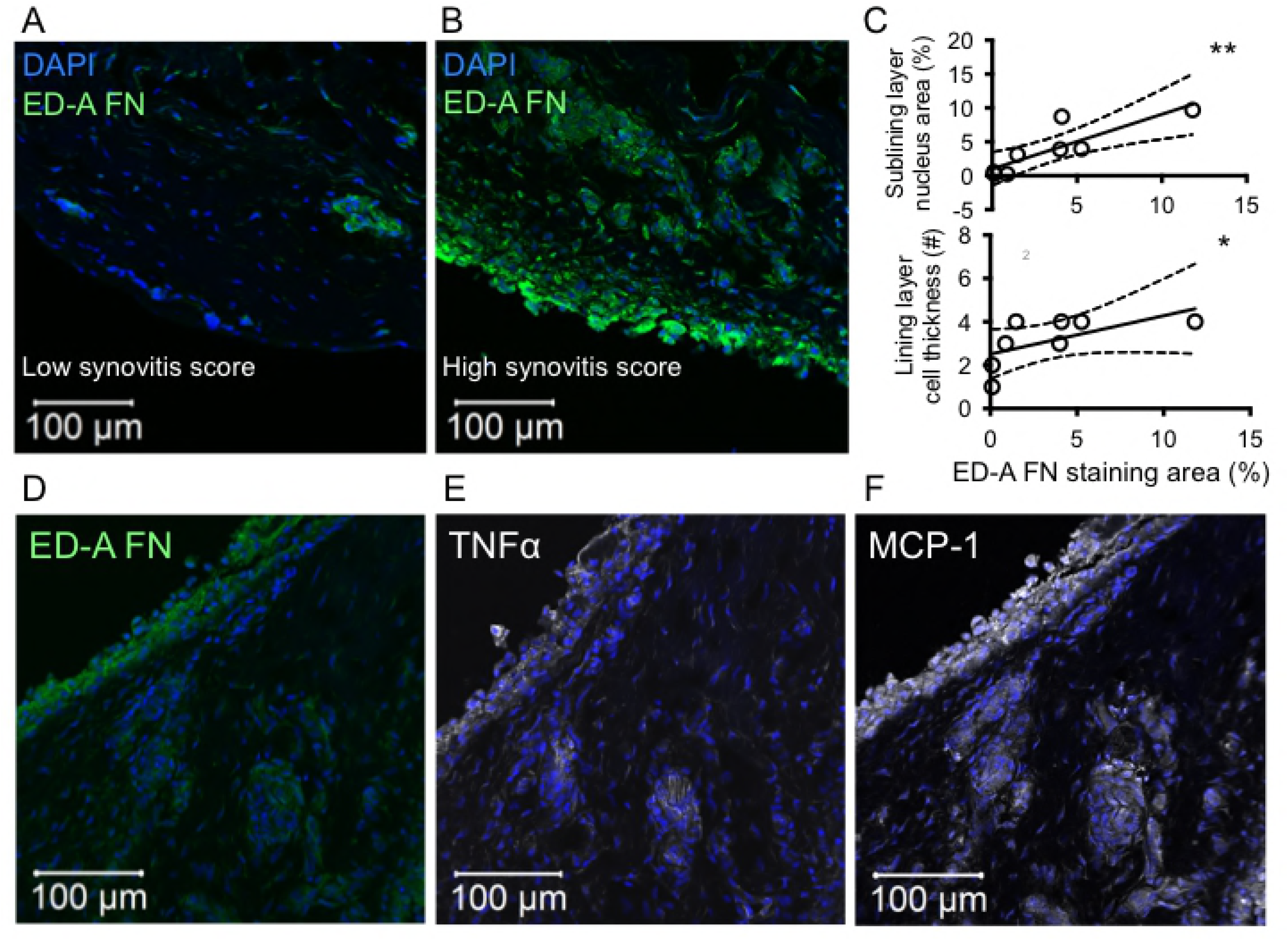
ED-A FN expression and degree of synovitis in OA synovium and co- localization with MCP-1 and TNFα. **A and B**. Representative confocal microscopy images of ED-A FN staining and synovitis score (n=8). **C**. ED-A FN expression associated with cell infiltration in the sublining layer and cell thickness of the lining layer. Data were analyzed using the Spearman Rho. **D-F**. Representative confocal microscopy images of localization of ED-A FN, TNFα and MCP-1 in OA synovium (n=5). TNFα and MCP-1 staining was found in areas with ED-A FN staining. * *P*<0.05, ** *P*<0.01.

### ED-A FN is located to areas of increased MCP-1 and TNFα in OA synovium

To test whether this association was due to expression of pro-inflammatory molecules we stained the OA synovial membranes with both ED-A FN MCP-1 and TNFα. ED-A FN and MCP-1 were co-stained on the same slides using antibodies with different fluorophores while TNFα was stained on serial slides. ED-A FN co-localized with both MCP-1 and TNFα in all stained synovial membranes (n=5) (Figure 2D-F and Supplementary figure 3). The staining of MCP-1 and TNFα was mostly located to cells in close proximity the ED-A FN positive cells but not specifically to the ED-A FN positive cells.

### Recombinant ED-A FN increase the secretion of MCP-1 and TNFα by monocyte-derived macrophages and OA synovial cells

We now speculated that the association between ED-A FN and MCP-1 and TNFα in the synovial membrane could be caused by a stimulatory effect of ED-A FN on MCP-1 and TNFα production by synovial macrophages. Monocyte-derived macrophages and OA synovial cells were therefore incubated with recombinant ED-A FN. Recombinant ED-A FN increased the production of TNFα from RAW 264.7 cells using a concentration of 10 μg/ml (n=3) (Figure 3A). This stimulatory effect was reduced when treating the ED-A FN with proteinase K or heat but not with polymyxin B (n=3) (Figure 3A). Recombinant ED-A FN further increased the production of MCP-1 significantly in monocyte-derived macrophages and increased both MCP-1 and TNFα by both monocyte-derived macrophages (n=3) and OA synovial cells (n=3) in all cultures tested (both MCP-1 and TNFα concentrations were below the detection limit of the ELISA assays in one culture) (Figure 3B and C). ED-A FN was used at a rather low concentration of 100 ng/ml for direct comparison with LPS and the stimulatory capacity was lower compared with LPS. These findings indicate that ED-A FN has the capacity to induce the production of pro-inflammatory molecules in OA synovial macrophages.

**Figure 3.**
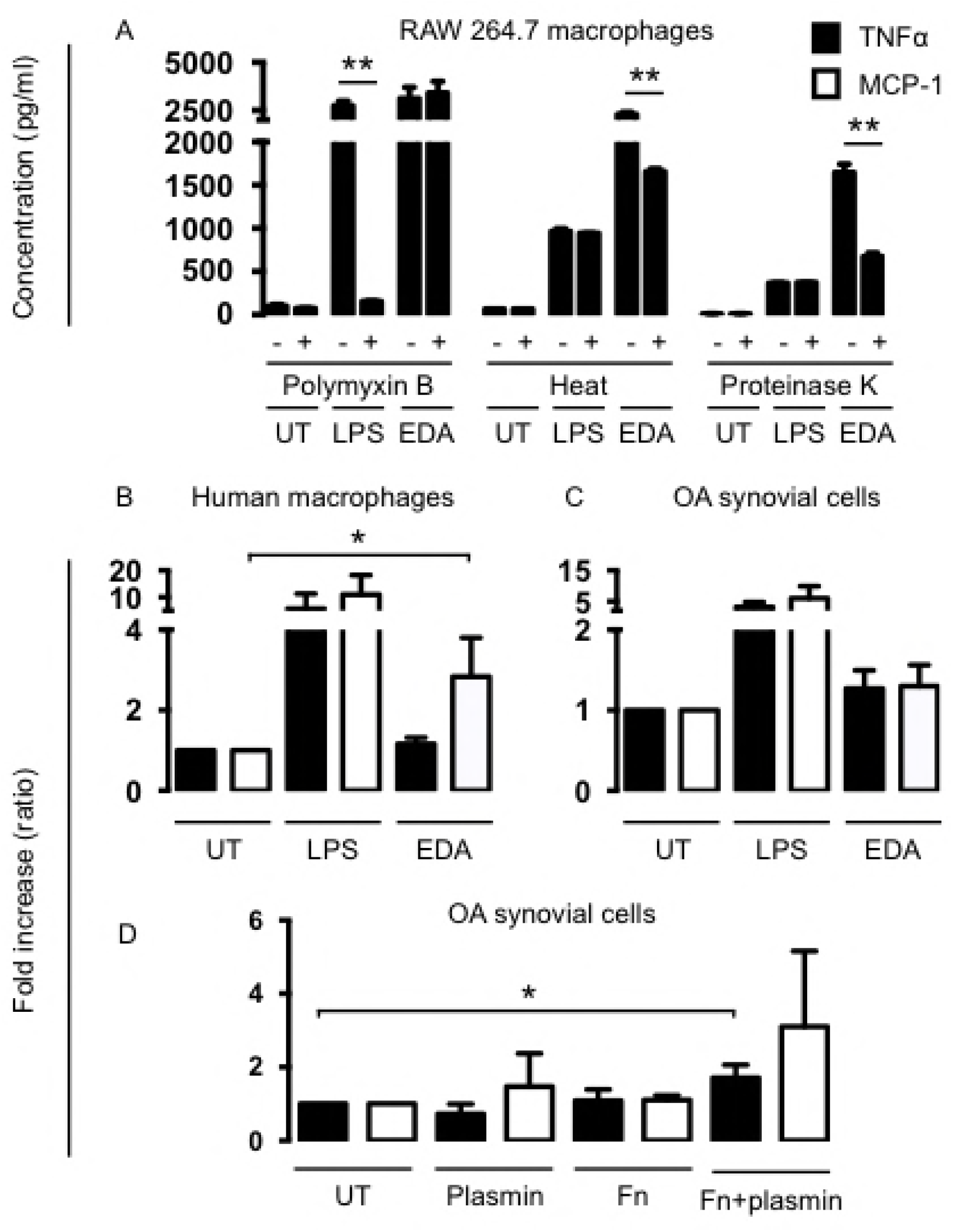
Effect of ED-A FN and plasmin derived FN fragments on TNFα and MCP-1 production. **A**. RAW 264.7 cells incubated untreated (UT), with LPS (1 ng/ml), or with a supraphysiological concentration of recombinant ED-A FN (10 μg/ml) (n=3). **B**. Human macrophages incubated UT, with LPS (100 ng/ml), or with a physiological concentration of recombinant ED-A FN (100 ng/ml) (n=3). **C**. OA synovial cell cultures incubated UT, with LPS (100 ng/ml), or recombinant ED-A FN (100 ng/ml) (n=3). **D**. OA synovial cell cultures incubated with medium, plasmin (0.5 ng/ml), cellular FN (1 μg/ml), or FN fragments generated by incubating cellular FN (1 μg/ml) with plasmin (0.5 ng/ml) (n=3). Ratios were made by dividing the concentration in supernatants from treated cells with the concentration in supernatants from untreated cells. Data were log transformed and analyzed with the paired t-test. Bars indicate the median and whiskers indicate the IQR. * *P*<0.05. ** *P*<0.01.

### Fibronectin exposed to plasmin increases the secretion of MCP-1 and TNFα by OA synovial cells

Finally, we wanted to demonstrate that the FN fragments containing the ED-A domain are able to generate the same stimulatory response as recombinant ED-A FN. Cellular FN containing the ED-A domain was therefore incubated with plasmin to generate fragments. These FN fragments were then added to OA synovial cells. The ED-A containing FN fragments increased the production of both MCP-1 and TNFα from the OA synovial cells in all cultures (n=3) (Figure 3D).

## Discussion

OA is a common joint disease involving wear and tear and low-grade inflammation. The disease pathogenesis has thus been speculated to resemble a pathogenic wound healing response [1,2]. Here, we show that ED-A FN could be part of such pathogenic wound healing mechanisms as we demonstrate that TGF? stimulates myofibroblast production of ED-A FN, and that ED-A FN fragments stimulate macrophage production of pro-inflammatory mediators.

First, we studied potential inducers of ED-A FN secretion in OA. We found formation of ED-A FN in αSMA positive myofibroblasts isolated from OA synovial tissue and co- localization of ED-A FN and αSMA in the OA synovium. This is in line with previous findings suggesting that ED-A FN is expressed in TGFβ induced myofibroblast differentiation [6]. Further, the formation of ED-A FN was increased by both OA synovial fluid and a protein suspension of lysed chondrocytes. This could be important in relation to OA because synovial fluid resembles the local milieu in the OA joint and the lysed chondrocytes likely contains molecules released by wear and tear and cartilage damage. Our findings could in part be explained by previous findings of increased TGFβ in OA synovial fluid [23] and increased FN secretion after blunt cartilage trauma in a bovine model of OA [24]. Further, the role of a general wound healing response in OA is supported by findings of increased fibrosis and joint stiffening in OA [25], and by the protective effect of blocking TGFβ in experimental OA in mice [4]. However, it is not known whether the induction of ED-A FN by OA synovial fluid and lysed chondrocytes in this study is a result of the presence of proteins, nucleic acids, lipids or other components.

Then, we studied the effects of ED-A FN in OA. We found that ED-A FN is associated with lining layer thickness and sublining cellular density in the OA synovium. This is in line with previous findings of high ED-A FN expression in rheumatoid arthritis and in proliferative regions of OA [26,27].

A recent study however, only found small amounts of ED-A FN mRNA in OA synovium [28]. This suggests that the formation of ED-A FN is rather a result of enzymatic digestion or folding than synthesis. The association between ED-A FN expression and cellular density and presence of inflammatory molecules in the OA synovium is of particular interest because there are currently ongoing studies of utilizing ED-A FN as a drug delivery target in rheumatoid arthritis and other diseases [29]. E.g., Dekavil is a drug combining a human F8 antibody specific to the ED-A domain of FN fused to the anti-inflammatory cytokine IL-10 [30,31].

Further, ED-A FN staining was located in close proximity to staining of MCP-1 and TNFα. This could in part explain the association between ED-A FN expression and increased lining layer thickness and cellular infiltrate. Macrophages are known to be part of the cellular infiltrate in the OA synovium [32]. We therefore studied the effect of recombinant ED-A FN on macrophage production of these pro-inflammatory mediators.Recombinant ED-A FN stimulated the production of MCP-1 and TNFα by monocyte- derived macrophages and synovial cells. This supports previous findings of ED-A FN induced increase of IL-1β production by synovial cells [33]. The stimulatory effects could be due to ED-A FN binding to TLR-4 as previously reported [14].

Finally, we studied the potential generation of FN fragments in OA. FN fragments generated by pre-incubating cellular FN with plasmin stimulated the secretion of MCP-1 and TNFα by monocyte-derived macrophages and synovial cells. Our data support that FN fragments containing ED-A are generated in OA as a consequence of proteolytic activity as proposed previously [34]. These fragments would be able to stimulate TLR4. However, other mechanisms for FN fragment stimulation of the synovial cells cannot be excluded. E.g., it was recently shown that FN fragments also have the potential to activate TLR2 [35], and that FN fragments not containing ED-A bind TLR4 [36].

There is an unmet therapeutic need preventing OA disease progression. This study supports that ED-A FN could be a target for drug delivery to the inflamed joint or that the formation of ED-A FN or the enzymatic fragmentation of FN could be novel therapeutic targets in OA.

## Conclusion

Taken together, we show that myofibroblasts produce ED-A FN during wound healing-like conditions in OA. Fragments containing the ED-A domain stimulate MCP-1 and TNFα production by macrophages possibly promoting inflammation in OA.

## List of abbreviations

αSMA: alpha smooth muscle actin
DAMPs: danger associated molecular patterns ECM
ED-A FN: fibronectin containing the ED-A domain
FBS: fetal bovine serum
FLS: fibroblast-like synovial cells
Ig: immunoglobulin
IL: interleukin
IQR: interquartile range
LPS: lipopolysaccharide
MCP-1: monocyte chemoattractant protein 1
MDM: monocyte-derived macrophages
OA: osteoarthritis
PBMC: peripheral blood mononuclear cell
PBS: phosphate buffered saline
TGFβ: tissue growth factor beta
TLR: Toll-like receptor
TNFα: tumour necrosis factor alpha
UT: untreated

## Declarations

### Ethical Approval and Consent to participate

All study participants gave ethical approval and consent to participate.

### Consent for publication

All study participants gave consent for publication.

### Availability of supporting data

Please contact author for data requests.

### Competing interests

The authors have no conflicts of interest.

### Funding

This work was supported by the Danish Research council and the Faculty of Health at Aarhus University.

## Authors’ contributions

All authors were involved in drafting the article or revising it critically for important intellectual content, and all authors approved the final version to be published.Study conception and design: Kragstrup, Robinson, Sokolove. Acquisition of data: Kragstrup, Sohn, Onuma, Lepus, Wang, Sokolove. Analysis and interpretation of data: Kragstrup, Sohn, Robinson, Sokolove All authors read and approved the final manuscript.

## Authors’ information

**Supplementary figure 1. ED-A FN production by OA FLSs**. Representative confocal microscopy images of αSMA (red) and ED-A FN (green) in OA FLS cultures incubated with medium.

**Supplementary figure 2. ED-A FN expression in OA synovium.** Representative confocal microscopy images of ED-A FN (green) co-localization with αSMA (red) (n=3).

**Supplementary figure 3. ED-A FN co-localization with MCP-1 and TNFα**. Representative confocal microscopy images of localization of ED-A FN, TNFα and MCP-1 in OA synovium (n=5).

## References

1. Scanzello CR, Plaas A, Crow MK. Innate immune system activation in osteoarthritis: is osteoarthritis a chronic wound? Curr Opin Rheumatol. 2008;20:565–72.

2. Sokolove J, Lepus CM. Role of inflammation in the pathogenesis of osteoarthritis: latest findings and interpretations. Therapeutic Advances in Musculoskeletal Disease. 2013.

3. Robinson WH, Lepus CM, Wang Q, Raghu H, Mao R, Lindstrom TM, et al. Low-grade inflammation as a key mediator of the pathogenesis of osteoarthritis. Nat Rev Rheumatol. 2016;12:580–92.

4. Scharstuhl A, Vitters EL, van der Kraan PM, van den Berg WB. Reduction of osteophyte formation and synovial thickening by adenoviral overexpression of transforming growth factor β/bone morphogenetic protein inhibitors during experimental osteoarthritis. Arthritis Rheum. 2003;48:3442–51.

5. Erridge C. Endogenous ligands of TLR2 and TLR4: agonists or assistants? J. Leukoc. Biol. 2010;87:989–99.

6. White ES, Muro AF. Fibronectin splice variants: Understanding their multiple roles in health and disease using engineered mouse models. IUBMB Life. 2011;63:538–46.

7. Gonzales-Gronow M, Enghild JJ, Pizzo SV. Streptokinase and human fibronectin share a common epitope: implications for regulations of fibrinolysis and rheumatoid arthritis. Biochimica et Biophysica Acta (BBA) - Molecular Basis of Disease. 1993;1180:283–8.

8. Peters JH, Loredo GA, Benton HP. Is osteoarthritis a ‘fibronectin-integrin imbalance disorder’? Osteoarthr. Cartil. 2002;10:831–5.

9. Barilla ML, Carsons SE. Fibronectin fragments and their role in inflammatory arthritis. Semin. Arthritis Rheum. 2000;29:252–65.

10. Homandberg GA, Meyers R, Williams JM. Intraarticular injection of fibronectin fragments causes severe depletion of cartilage proteoglycans in vivo. J. Rheumatol. 1993;20:1378–82.

11. Xie DL, Meyers R, Homandberg GA. Fibronectin fragments in osteoarthritic synovial fluid. J. Rheumatol. 1992;19:1448–52.

12. Homandberg GA, Hui F. Association of proteoglycan degradation with catabolic cytokine and stromelysin release from cartilage cultured with fibronectin fragments. Arch. Biochem. Biophys. 1996;334:325–31.

13. Carsons S. Extra domain-positive fibronectins in arthritis: wolf in sheep’s clothing? Rheumatology (Oxford). 2001;40:721–3.

14. Okamura Y, Watari M, Jerud ES, Young DW, Ishizaka ST, Rose J, et al. The extra domain A of fibronectin activates Toll-like receptor 4. J. Biol. Chem. 2001;276:10229–33.

15. Przybysz M, Borysewicz K, Katnik-Prastowska I. Differences between the early and advanced stages of rheumatoid arthritis in the expression of EDA-containing fibronectin. Rheumatol. Int. 2009;29:1397–401.

16. Shiozawa K, Hino K, Shiozawa S. Alternatively spliced EDA-containing fibronectin in synovial fluid as a predictor of rheumatoid joint destruction. Rheumatology (Oxford). 2001;40:739–42.

17. Muro AF, Chauhan AK, Gajovic S, Iaconcig A, Porro F, Stanta G, et al. Regulated splicing of the fibronectin EDA exon is essential for proper skin wound healing and normal lifespan. J Cell Biol. Rockefeller Univ Press; 2003;162:149–60.

18. Ffrench-Constant C, Van de Water L, Dvorak HF, Hynes RO. Reappearance of an embryonic pattern of fibronectin splicing during wound healing in the adult rat. J Cell Biol. 1989;109:903–14.

19. Serini G, Bochaton-Piallat ML, Ropraz P, Geinoz A, Borsi L, Zardi L, et al. The fibronectin domain ED-A is crucial for myofibroblastic phenotype induction by transforming growth factor-beta1. J Cell Biol. 1998;142:873–81.

20. Kellgren JH, Lawrence JS. Radiological assessment of osteo-arthrosis. Ann. Rheum. Dis. BMJ Group; 1957;16:494–502.

21. Altman R, Asch E, Bloch D, Bole G, Borenstein D, Brandt K, et al. Development of criteria for the classification and reporting of osteoarthritis. Classification of osteoarthritis of the knee. Diagnostic and Therapeutic Criteria Committee of the American Rheumatism Association. Arthritis Rheum. 1986. pp. 1039–49.

22. Kragstrup TW, Jalilian B, Hvid M, Kjærgaard A, Østgård R, Schiøttz Christensen B, et al. Decreased plasma levels of soluble CD18 link leukocyte infiltration with disease activity in spondyloarthritis. Arthritis Res. Ther. 2014;16:R42.

23. Schlaak JF, Pfers I, Meyer Zum Büschenfelde KH, Märker-Hermann E. Different cytokine profiles in the synovial fluid of patients with osteoarthritis, rheumatoid arthritis and seronegative spondylarthropathies. Clin. Exp. Rheumatol. 1996;14:155–62.

24. Ding L, Guo D, Homandberg GA, Buckwalter JA, Martin JA. A single blunt impact on cartilage promotes fibronectin fragmentation and upregulates cartilage degrading stromelysin-1/matrix metalloproteinase-3 in a bovine ex vivo model. J. Orthop. Res. 2014;32:811–8.

25. Hill CL, Hunter DJ, Niu J, Clancy M, Guermazi A, Genant H, et al. Synovitis detected on magnetic resonance imaging and its relation to pain and cartilage loss in knee osteoarthritis. Ann. Rheum. Dis. 2007;66:1599–603.

26. Hino K, Shiozawa S, Kuroki Y, Ishikawa H, Shiozawa K, Sekiguchi K, et al. Eda-containing fibronectin is synthesized from rheumatoid synovial fibroblast-like cells. Arthritis Rheum. 1995;38:678–83.

27. Kriegsmann J, Berndt A, Hansen T, Borsi L, Zardi L, Bräuer R, et al. Expression of fibronectin splice variants and oncofetal glycosylated fibronectin in the synovial membranes of patients with rheumatoid arthritis and osteoarthritis. Rheumatol. Int. 2004;24:25–33.

28. Scanzello CR, Markova DZ, Chee A, Xiu Y, Adams SL, Anderson G, et al. Fibronectin splice variation in human knee cartilage, meniscus and synovial membrane: observations in osteoarthritic knee. J. Orthop. Res. 2015;33:556–62.

29. Kumra H, Reinhardt DP. Fibronectin-targeted drug delivery in cancer. Adv. Drug Deliv. Rev. 2016;97:101–10.

30. Schwager K, Kaspar M, Bootz F, Marcolongo R, Paresce E, Neri D, et al. Preclinical characterization of DEKAVIL (F8-IL10), a novel clinical-stage immunocytokine which inhibits the progression of collagen-induced arthritis. Arthritis Res. Ther. 2009;11:R142.

31. Galeazzi M, Bazzichi L, Sebastiani GD, Neri D, Garcia E, Ravenni N, et al. A phase IB clinical trial with Dekavil (F8-IL10), an immunoregulatory “armed antibody” for the treatment of rheumatoid arthritis, used in combination wiIh methotrexate. Isr. Med. Assoc. J. 2014;16:666.

32. Raghu H, Lepus CM, Wang Q, Wong HH, Lingampalli N, Oliviero F, et al. CCL2/CCR2, but not CCL5/CCR5, mediates monocyte recruitment, inflammation and cartilage destruction in osteoarthritis. Ann. Rheum. Dis. 2017;76:914–22.

33. Saito S, Yamaji N, Yasunaga K, Saito T, Matsumoto S, Katoh M, et al. The fibronectin extra domain A activates matrix metalloproteinase gene expression by an interleukin-1- dependent mechanism. J. Biol. Chem. 1999;274:30756–63.

34. Peters JH, Carsons S, Kalunian K, McDougall S, Yoshida M, Ko F, et al. Preferential recognition of a fragment species of osteoarthritic synovial fluid fibronectin by antibodies to the alternatively spliced EIIIA segment. Arthritis Rheum. 2001;44:2572–85.

35. Hwang HS, Park SJ, Cheon EJ, Lee MH, Kim HA. Fibronectin fragment-induced expression of matrix metalloproteinases is mediated by MyD88-dependent TLR-2 signaling pathway in human chondrocytes. Arthritis Res. Ther. 2015;17:320.

36. Sofat N, Robertson SD, Wait R. Fibronectin III 13-14 domains induce joint damage via Toll-like receptor 4 activation and synergize with interleukin-1 and tumour necrosis factor. J Innate Immun. 2012;4:69–79.

